# Effects of crude oil on juvenile threespine stickleback somatic and immune development

**DOI:** 10.1101/2021.11.17.469020

**Authors:** Kelly S. Ireland, Kathryn Milligan-Myhre

## Abstract

Aquatic oil spills have resounding effects on surrounding ecosystems, and thus significant resources are committed to oil spill responses to remove the oil from the environment as quickly as possible. Oil has immunotoxic effects and may be particularly harmful to larval and juvenile fish as it can cause a number of developmental defects and stunt growth. In spite of significant efforts to clean oil, it is unclear whether larval and juvenile fish can recover from the effects of oil and no work has been done on the effect crude oil has on developing threespine stickleback (Gasterosteus aculeatus) fish. Threespine stickleback are a ubiquitous sentinel species in the northern hemisphere and are an important food source for many larger, economically valuable fish. As fish with fully marine, anadromous, and freshwater populations, stickleback are exposed to oil in a variety of aquatic environments. We hypothesized that oil exposure would suppress both growth and immunity of developing stickleback, but that fish health could be recovered by removal of the crude oil. Fish were exposed to Alaska North Slope crude oil and then were moved to water without crude oil for two weeks (depuration). Measurements of growth and immunity were taken before and after the depuration. We found that crude oil effected different developmental pathways independently, significantly impacting some but not others. This is the first study to examine the effect crude oil has on early stages of stickleback development, and that stickleback fish are unable to recover from exposure after being transferred to clean water for two-weeks, suggesting larval/juvenile stickleback exposed to crude oil need longer than two-weeks to recover if they are able to recover at all.

## 1. Introduction

Thousands of oil spills occur annually around the world. Crude oil has profound impacts on fish species, including alteration of reproduction, mortality, cardiac function, development, immunity, and the microbiota (Bayha et al., 2017; Heintz et al., 2000; Incardona et al., 2004; Benjamin J Laurel et al., 2019; Reynaud and Deschaux, 2006; Song et al., 2011). Crude oil and polycyclic aromatic hydrocarbons (PAHs) found in crude oil also suppress immunity in fish, such as southern flounder, Japanese flounder, and Chinook salmon (Arkoosh and Collier, 2002; Bayha et al., 2017; Song et al., 2011). For example, crude oil increases susceptibility to the pathogenic bacteria Vibrio anguillarum in southern flounder (Paralichthys lethostigma) and Chinook salmon (Oncorhynchus tshawytscha) by suppressing the immune system, which leads to increased mortality (Arkoosh et al., 2002; Arkoosh and Collier, 2002; Bayha et al., 2017). Exposure to crude oil has acute and chronic effects on salmon fisheries, such as decreased marine survival and decreased reproductive output, which can impact runs for many salmon generations (Heintz et al., 2000).

Oil exposure during development is arguably the worst time for fish to be exposed as crude oil can cause many developmental defects during this time (Heintz et al., 2000; Incardona et al., 2004; Benjamin J Laurel et al., 2019). Embryonic crude oil exposure in Atlantic haddock (Melanogrammus aeglefinus) and polar cod (Boreogadus saida) causes cardiac and craniofacial defects (Benjamin J. Laurel et al., 2019; Sørhus et al., 2016). PAH exposure in embryonic zebrafish (Danio rerio) causes spinal curvature (Incardona et al., 2004). Pink salmon exposed to oil as juveniles have decreased marine survival (Heintz et al., 2000). These defects can affect a fish’s ability to survive and, thus, overall population survival.

Recovery from crude oil exposure after oil is removed, and fish are allowed to depurate, is species, time, and tissue dependent. For example, juvenile pink salmon exposed to crude oil weighed less than those fish not exposed, but at maturity (after oil is removed) there were no differences, although not as many oil treated fish survived to maturity (Heintz et al., 2000). In Australian Crimson-Spotted Rainbowfish crude oil water accommodated fractions (WAFs) caused an increase in citrate synthase, lactate dehydrogenase, and ethoxy-resorufin-O-deethylase (EROD), but all levels returned to normal after a two-week depuration period (Pollino and Holdway, 2003). In Seahorse Hippocampus reidi exposed to crude oil, genotoxic damage was increased (number of micronuclei), but recovery was possible after 7 days (Delunardo et al., 2013). Transcription of most genes in rainbow trout exposed to crude oil returned to normal after a 96 hour recovery period in gill tissue, but differential transcription was still observed in liver tissue after the recovery period, suggesting that recovery can be tissue dependent (Hook et al., 2010). Immune suppression following exposure to heavy crude oil is observed in Japanese flounder, but fish were able to recover after about 1 week (Song et al., 2012b). Unfortunately, most studies with a recovery time period did not use statistical analysis that allowed for the examination of the interaction between treatment and time, but rather the effect of treatment and time individually or just the effect of treatment at each sampling time point.

Threespine stickleback fish (Gasterosteus aculeatus) are important to the ecosystem as a ubiquitous sentinel species and food source in the northern hemisphere, existing as anadromous, oceanic, and freshwater populations (Gard and Bottorff, 2014). Decreased stickleback populations due to crude oil spills can limit food sources for other species already challenged with oil exposure. Anadromous stickleback can be exposed to marine oil spills as adults in the ocean or as spawning adults, embryos, larvae, and juveniles in estuaries and streams if oil spreads into these waterways. Freshwater populations of stickleback can be exposed to oil from oil pipeline spills or spills from other ground transport or storage of crude oil. Stickleback eggs do not float, so eggs would not be exposed to surface slicks, but rather dispersed oil in the water column or oil that sinks to the bottom. Stickleback eggs are smaller (1.33-2.16 mm) than fish species typically used for studies on the effects of crude oil or PAHs, such as salmon (6 mm) (Glippa et al., 2017; Thorn and Morbey, 2018). Fish eggs that are large and have a smaller surface area to volume ratio do not as rapidly accumulate PAHs as smaller eggs, so stickleback embryos may be more susceptible to the effects of crude oil (Edmunds et al., 2015).

The objective of this study was to determine how developing stickleback are affected by crude oil exposure, specifically how growth and the expression of two immune genes would be affected, and if fish are able to recover after crude oil removal.

Statistical analysis that addresses the interaction between treatment and time was used for this study, making inferences about recovery more robust than some previous studies. Fish were exposed to Alaska North Slope crude oil for one-week, between 7- and 14-days post fertilization (dpf), followed by a two-week depuration period. Somatic measurements and immune gene expression were measured at both 14 and 28 dpf. This experiment was the first to characterize morphological and immune impacts and resiliency of threespine stickleback to crude oil and the first to use larval/juvenile stage threespine stickleback in an oil toxicology study.

## 2. Materials and Methods

### 2.1 Fish Husbandry

Adult fish were collected from Bear Paw Lake in Houston, Alaska (61.61396, −149.75309) under Alaska Department of Fish and Game permit P-18-006 on October 5, 2018. These fish were acclimated in the lab for four months under a short light cycle of 10 hours of light per day, and then shifted to 20 hours of light per day to induce egg and viable sperm production. Embryos were generated from in vitro crosses on February 7, 2019. Sperm from 2 males were mixed and used to fertilize egg clutches from 7 females, following an IACUC approved protocol (IACUC #1302464-1, 1302466-3). Eggs were fertilized, incubated for several hours to ensure viability, and then viable eggs were distributed randomly to six culture flasks (47 fish/flask) with filter caps (150 cm2, TPP Techno Plastic Products AG, Trasadingen, Switzerland), containing 175 ml of 4 ppt instant ocean stickleback embryo medium (SBEM). All fish were incubated at 18°C on a 20-hour light cycle in these flasks until 14 days post fertilization (dpf). At 8 dpf the flasks were flicked to release the remaining unhatched fish from their chorions. Randomly assigned flasks were exposed to oil at seven dpf while fish were still in the larval stage of development (details discussed in 2.3). Between 14 and 28 dpf the fish enter into the juvenile stage of development. At 14 dpf, fish not randomly selected for lethal sampling were moved into breeder nets within individual tanks for depuration on the same recirculating water that was free of crude oil, with an average salinity of 4.16 ppm and kept on a 20-hour light cycle at 16.3-17.6°C. Fish were fed brine shrimp and Golden Pearl Reef and Larval Fish flake food (Brine Shrimp Direct, Ogden, Utah, USA) daily. Mortality was recorded daily.

### 2.2 Image and Tissue Collection

At 14 and 28 dpf, a subset of each flask or tank respectively (n=12/flask or tank, except for one tank, n=8, where only 8 fish survived to 28 dpf) were euthanized using a lethal dose of buffered MS-222 (IA-CUC #1363107-1). Standardized images of each fish were taken. After removing intestines for future sequencing analysis (DNA yield was deemed too low for analysis so no data is available), tissues from three individuals from the same tank were added to 200 µl Trizol with a 1:1 mix of 0.5 and 1.0 mm zirconia oxide beads in a RINO tube (Next Advance Inc. Troy, NY, USA), flash-frozen using liquid nitrogen, and stored at −80°C until processing. A timeline of the experiment and sample processing workflow is detailed in Fig. 1.

**Fig. 1.**
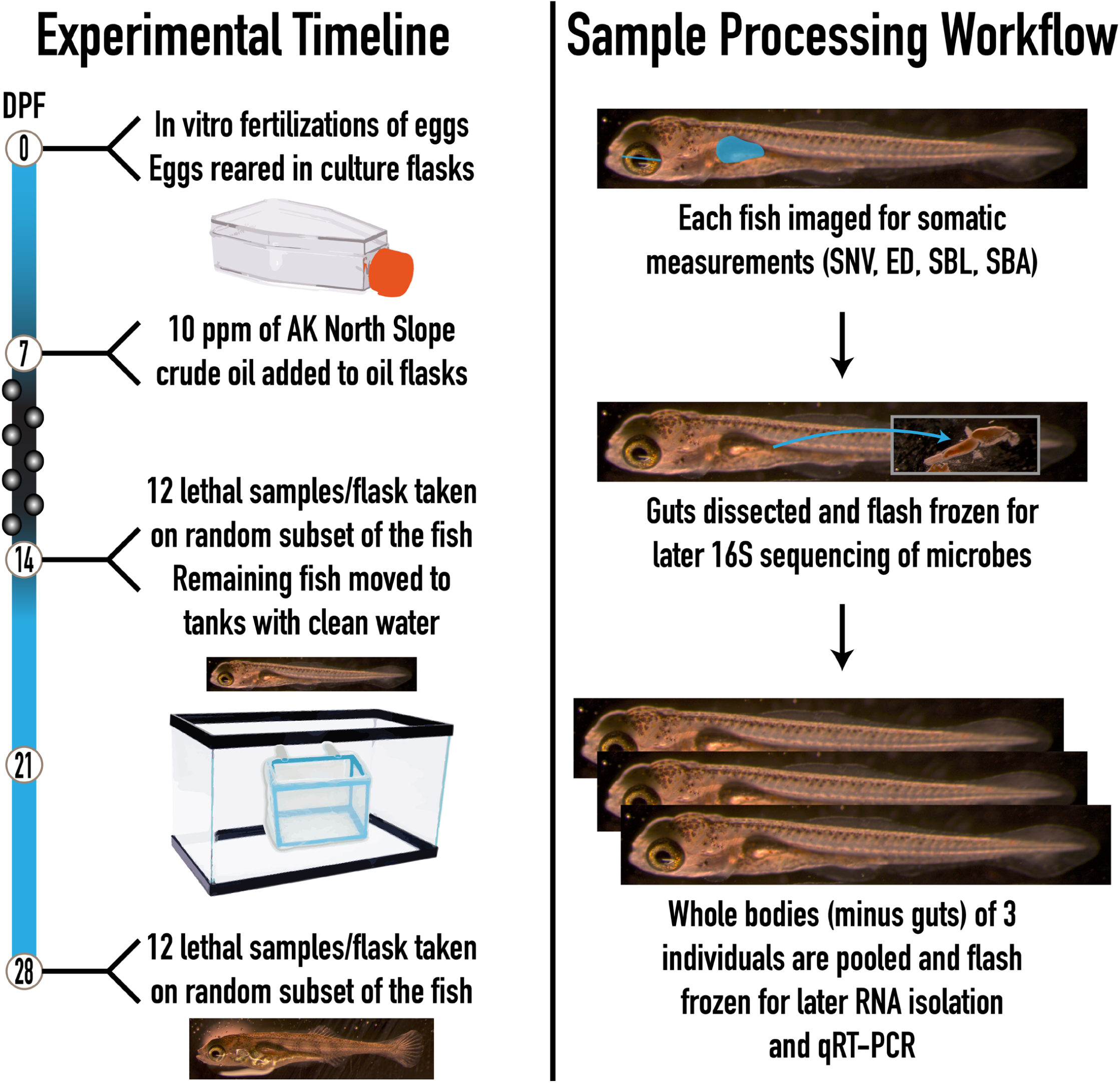
Experimental Timeline and Workflow. This workflow was used to determine the effects of crude oil on juvenile threespine stickleback. Fish were reared in culture flasks from 1-14 dpf and in tanks from 14-28 dpf. Alaska North Slope crude oil or untreated water was added to randomly assigned flasks at seven dpf. Samples of 12 fish/flask were taken at 14 and 28 dpf. Fish were imaged, guts were dissected out (DNA yield too low for any analysis), and the whole bodies of three fish (from the same flask) were pooled for RNA isolation and qRT-PCR.

### 2.3 Treatments

To determine the effects of crude oil on threespine stickleback, randomly assigned flasks were exposed to either 10 mL of parental water containing no crude oil (3 flasks) or 10 mL of a crude oil/water mixture (3 flasks) at seven dpf (IACUC #1363107-1), prior to addition, 10 mL of SBEM were removed from each flask. The oil mixture of Alaska North Slope Crude Oil was prepared by adding 1.2 µL of crude oil to 10 mL of SBEM in a 15 mL falcon tube and then shaking the crude oil and water for 11.5 hours on a benchtop shaker at room temperature before addition to the flasks. Fish remained in these flasks, with no additions of water or water changes, until fish were moved to tanks with freshwater at 14 dpf. An equal volume of parental populations’ water was added at seven dpf to the three control flasks and were moved to tanks at 14 dpf. Samples and data were collected at both 14 and 28 dpf.

### 2.4 Somatic Development

Somatic development was measured using ImageJ software on images taken directly before gut dissections at 14 and 28 dpf on a dissecting scope. All photos from the same collection were taken with the same magnification, along with an image of a ruler for scale. Snout-vent length, eye diameter, swim bladder length, and swim bladder area were all measured from each fish (Fig. 2). Each measurement was blinded for treatment and taken in triplicate by a single measurer and then averaged for analysis.

**Fig. 2.**
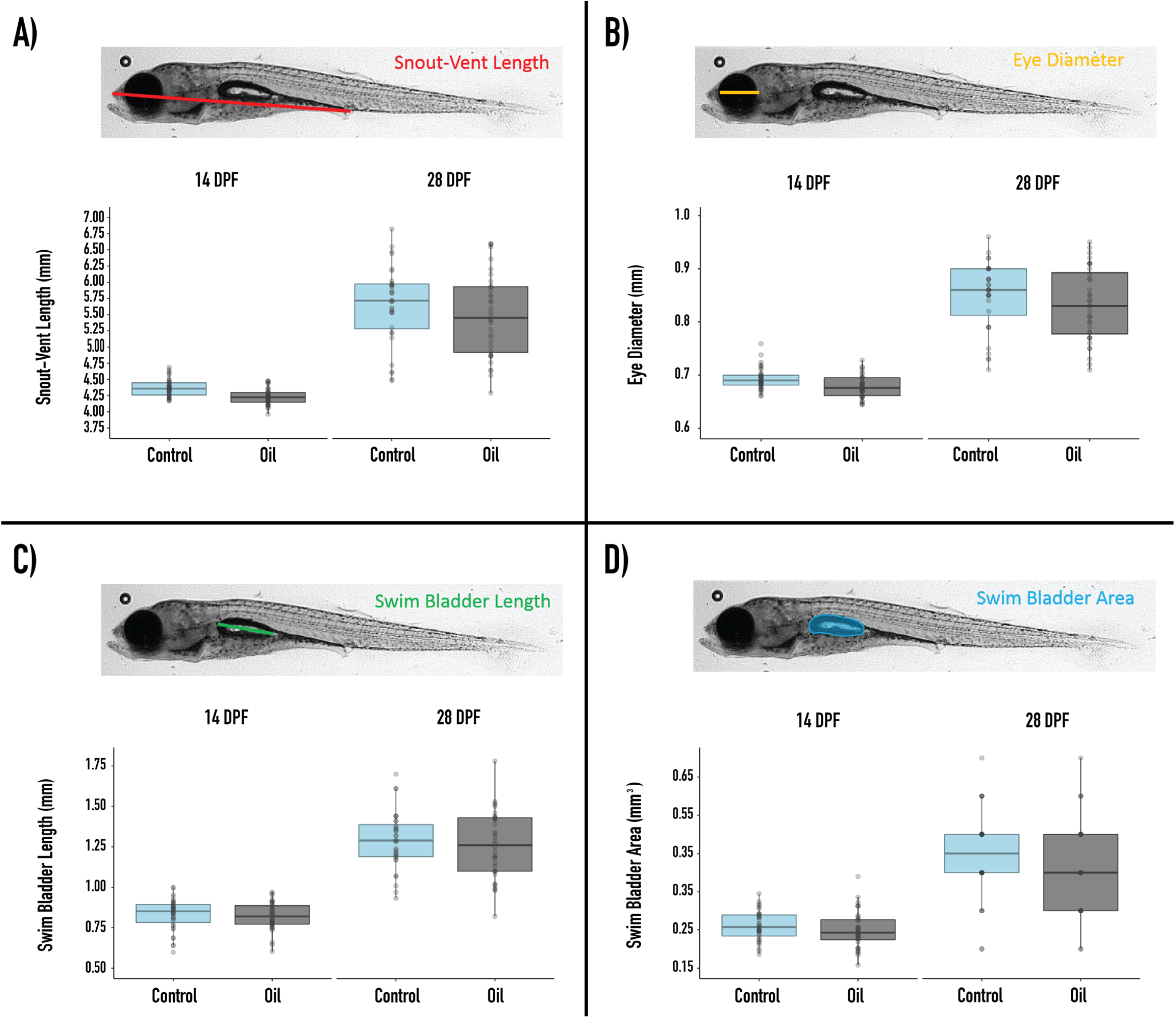
Somatic Measurements. Measurements of somatic growth in oil-treated and untreated fish. Snoutvent length **(A)**, eye diameter **(B)**, swim bladder length **(C)**, and swim bladder area **(D)** measurements were made on 12 fish from each flask (except one 28 dpf flask that only had eight fish survive) at 14 and 28 dpf. There were three flasks/treatment. Boxplots represent control (blue) and oil-treated (black) fish with the median, 25th, and 75th quartiles, whiskers, and all outlying data. Dots represent measurements of individual fish.

### 2.5 Immune Gene Expression

Expression levels of several innate immune system genes were measured using a well-established quantitative real-time polymerase chain reaction (qRT-PCR) protocol for samples taken at 14 and 28 dpf. A modified RNA isolation protocol using Trizol was used to isolate RNA from a pooled sample of 3 individuals (Leung and Dowling, 2005). Thawed samples were bead-beat on a Bead Mill24 homogenizer (Thermo Fisher Scientific, Waltham, Massachusetts, USA) before an additional 800 µl of Trizol was added, and the samples were flash-frozen again. The resulting homogenate was passed through a Qiashredder centrifuge column (Qiagen, Hilden, Germany) before two rounds of phase separation with chloroform in 2.0 phase lock gel tubes (QuantBio, Beverley, MA, USA) was performed. RNA was washed and eluted using the Qiagen RNAeasy Kit (Qiagen). RNA concentration, quality, and integrity were measured using a Qubit 4.0 Fluorometer, using the RNA High Sensitivity, Broad Range, and IQ Assay kits (Thermo Fisher Scientific). Average initial RNA concentration at 14 dpf was 67.11 ng/µl with a standard deviation of 28.59 ng/µl. Average initial RNA concentration at 28 dpf was 108.90 ng/µl with a standard deviation of 78.46 ng/µl. Initial RNA concentration did not significantly impact the expression of either gene. Working samples were created to a concentration of 50 ng/µl in RNase free water. Each sample was analyzed in triplicate qRT-PCR reactions using the Luna Universal One-Step RT-qPCR kit (New England Bio Labs, Ipswich, MA, USA) according to the manufacturer’s protocol. Each reaction contained 10 µl reaction mix, 1 µl WarmStart reverse transcriptase, 0.8 µl forward primer (10 µM), 0.8 µl reverse primer (10 µM), 6.4 µl RNase free water, and 1 µl standardized RNA sample (50 ng). Relative normalized expression of several innate immune genes that promote inflammation, interleukin 1 beta (IL-1ß), and tumor necrosis factor alpha (TNF-α) were analyzed using established immune gene primers for stickleback (Robertson et al., 2016). The qRT-PCRs were run on a BioRad CFX96 Touch Real-Time PCR Detection System (Bio-Rad Laboratories, Hercules, CA, USA). The BioRad CFX Manager software version 2.1 (Bio-Rad, 2013) calculated the relative normalized expression of IL-1ß and TNF-α against no template controls and published, well-established stickleback reference genes, specifically beta-2-microglobulin (B2M) and L13A ribosomal binding protein (RPL13A) (Hibbeler et al., 2008).

### 2.6 Statistical Analysis

The proportion of dead fish in control and oil treatment groups from 7-28 dpf (0-7 dpf was excluded as fish were not yet exposed to oil) was compared using an N-1 chi-squared test to assess differences in mortality. To assess the effects of oil exposure over time on both somatic responses and immune gene expression, a linear mixed-effects model of the following form was used:

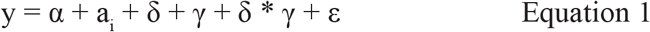

where y is the dependent variable (snout-vent length, eye diameter, swim bladder length, swim bladder area, IL-1ß or TNF-α), α is the overall intercept, ai is a random effect of flask i (1-6), δ is the effect of treatment (oil or control), γ is the effect of time (28 dpf or 14 dpf), δ * γ is the interaction of treatment and time, and ε is a residual with mean 0 and variance σ2. Because of apparent differences in variances between 14 and 28 dpf, the model estimated separate variances by time. When the interaction between treatment and dpf was not significant, a reduced model with the interaction term was removed, and only the results from the additive model were reported. Snout-vent length and swim bladder length could be allometrically correlated but were determined to not be correlated by a Pearson Correlation test (p= 0.926) and therefore were analyzed separately. Data and the R markdown file can be found at https://github.com/kellysue94/effects_of_crude_oil_on_juvenile_threespine_stickleback_020719.

## 3. Results

To determine how crude oil affects the survival, development, and immune system in developing stickleback, fish were exposed to Alaska North Slope crude oil between 7 and 14 dpf, and then were moved to tanks with water free of crude oil for a two-week depuration period. Somatic measurements and the expression of two immune genes were measured at both 14 and 28 dpf to determine how oil exposure affects growth and immunity and whether fish growth can recover after two weeks of depuration.

### 3.1 Mortality

Crude oil can increase mortality in fish, but this varies by concentration, species, and life stage in which fish were exposed. To determine whether crude oil increases mortality in developing stickleback, deaths of fish were recorded daily and mortality rates between 7-28 dpf were compared between the control and oil treatments. Oil exposure did not significantly affect mortality rates (p=0.9288, 5.08% mortality in control fish and 2.72% mortality in oil-treated fish).

### 3.2 Somatic Development

Snout-vent length, eye diameter, swim bladder length, and swim bladder area are all impacted by crude oil in other fish (Carls et al., 2000; Incardona et al., 2014, 2004; Benjamin J. Laurel et al., 2019; Li et al., 2019; Meador and Nahrgang, 2019; Raimondo et al., 2016; Sørhus et al., 2016). Thus, we measured each of these in stickleback one week after exposure (14 dpf) and after a two-week depuration (28 dpf). We hypothesized all somatic measurements would be reduced at both 14- and 28-days post fertilization (dpf) in fish exposed to crude oil compared to control treated fish. Dpf caused a significant increase in all somatic measurements (Fig. 2 and table 1) as the fish grew from 14 to 28 dpf (p=0.000 for all measurements). Both additive and interaction models were analyzed but given that the interaction term wasn’t significant, only the additive effect of dpf and treatment is reported. Oil exposure significantly reduced snout-vent length (Fig. 2A & Table 1, t=-4.14, df = 4, p=0.0144) and eye diameter (Fig. 2B, t=-2.86, df=4, p=0.0460) when time was held constant, meaning that snout-vent length and eye diameter were smaller in oil exposed fish across the course of the experiment, not just at 14 or 28 dpf. Oil exposure does not significantly change swim bladder length (Fig. 2C, t=0.00132, df=4, p=0.999) and swim bladder area (Fig. 2D, t=-0.500, p=0.644) over the course of the experiment. These results indicate that oil exposure significantly decreases snout-vent length and eye diameter, without recovery after a two-week depuration, but has no significant impact on swim bladder length or swim bladder area in this population of threespine stickleback, indicating that oil exposure effects development of some structures (like the length and eyes), but not others (like the swim bladder) at this concentration of crude oil.

### 3.3 Immune Gene Expression

Oil is immunotoxic to several fish species, but it is unknown whether oil exposure also causes immune suppression in stickleback. The expression of two innate immune genes involved in inflammation, interleukin 1 beta (IL-1ß) and tumor necrosis factor alpha (TNF-α), were measured. These innate immune genes were chosen as stickleback lack adaptive immunity until later in their juvenile stage and because the innate immunity appears to be more affected by oil exposure in other fish than adaptive immunity (Bo et al., 2012; Milligan-Myhre et al., 2016; Palm et al., 2003; Reynaud and Deschaux, 2006).

There was no significant effect of treatment alone across the course of the experiment on the expression of IL-1ß (t=2.490, df=4, p=0.1148) or TNF-α (t=1.442, df=4, p=0.2226) (Fig. 3). However, there was a significant positive effect of dpf on the expression of both genes (p=0.000) and a significant negative effect of the interaction between treatment and dpf in both IL-1ß (t=-2.543, df=39, p=0.0000) and TNF-α (t=-4.532, df=132, p=0.0000) indicating that in control fish IL-1ß and TNF-α tended to be upregulated over time (from 14 to 28 dpf). However, in oil-treated fish, there was a downward trend in the expression of both genes from 14 to 28 dpf. The variance in the expression of both genes is decreased in the oil treated group. A Bartlett test indicated that the variance of both IL-1ß expression (14 dpf – p= 0.000709,28 dpf - p=1.66e-05) and TNF-A expression (14 dpf – p=0.00133, 28 dpf – p=2.91e-06) in oil treated fish at 14 and 28 dpf was significantly reduced.

**Fig. 3.**
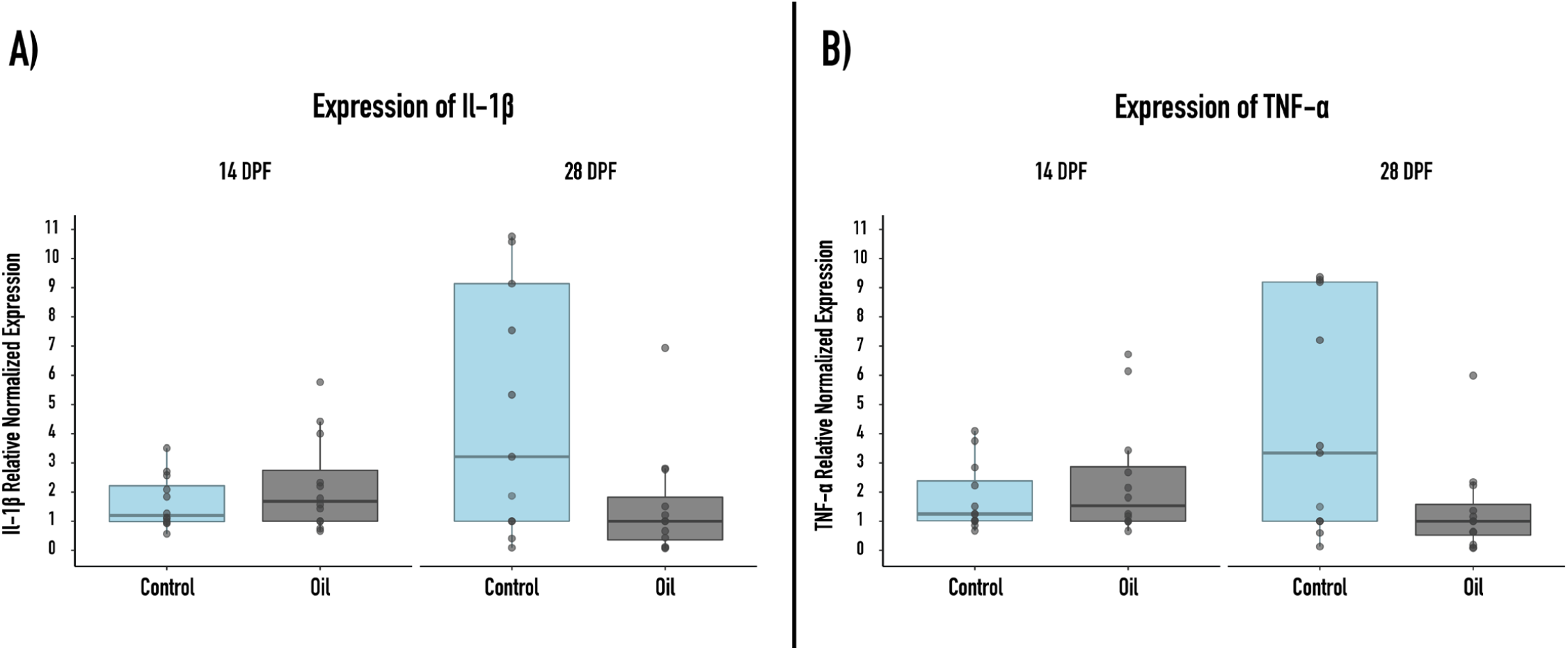
Immune Gene Expression. Measurements of innate immune genes in oil-treated and untreated fish. Relative normalized expression of IL-1ß **(A)** and TNF-α **(B)** was measured on four pooled samples of three individuals from each flask (except one 28 dpf flask that only had two pooled samples) at 14 and 28 dpf. There were three flasks/treatment. Boxplots represent control (blue) and oil-treated (black) fish with the median, 25th, and 75th quartiles, whiskers, and all outlying data. Dots represent individual measurements.

## 4. Discussion

This study is the first to examine how survival, growth, and innate immunity of developing threespine stickleback is affected by exposure to crude oil, and to determine whether the effects of crude oil in developing stickleback fish can be reversed by depuration.

### 4.1 Mortality

Oil exposure did not increase mortality in larval/juvenile threespine stickleback. In this experiment, the oil was added to flasks at seven dpf, just prior to hatching, which allowed fish to develop longer in their chorions before oil exposure, likely giving them an advantage in surviving. These results are similar to polar cod where mortality is not increased in fish exposed prior to hatching, but different than southern flounder where mortality is increased (Brown-Peterson et al., 2017, 2015; Benjamin J. Laurel et al., 2019). However, juvenile southern flounder and Liza ramada exposed to oil post-hatch do not have increased mortality (Bayha et al., 2017; Milinkovitch et al., 2011). In Japanese flounder post-hatch increases in mortality only occur at higher concentrations of crude oil at 0.3 g/L (Song et al., 2011), well above the exposure concentration in this study. Adult stickleback are also quite resilient to crude oil exposure, with a median tolerance of 6.89 mg/ liter (Moles et al., 1979), so it could be that developing stickleback are also fairly resilient to oil exposure. While oil may not cause any increases in mortality ex-situ, oil exposure could increase mortality in-situ. Oil induced decreases in snout-vent length and eye diameter could impair visual acuity and swim performance, which can impact prey availability and predation (Benjamin J Laurel et al., 2019). There is also a trend of downregulation of innate immune genes, IL-1ß and TNF-α, which may make fish more susceptible to pathogens, thus increasing mortality in the wild.

### 4.2 Somatic Development

Snout-vent length and eye diameter are both significantly decreased in developing threespine stickleback exposed to crude oil. In Sheepshead Minnow, there were significant differences in length and weight when exposed to PAHs at as low as 50 mg ∑PAH/kg dry sediment (Raimondo et al., 2016). In pink salmon, weight was lower in oil exposed fish at doses greater than 18 ppb before entering the marine environment, but there was no difference in size or weight when fish returned to their natal streams (Heintz et al., 2000). Crude oil exposure also decreases total length in Pacific herring (Carls et al., 2000). Eye diameter is also reduced by oil exposure in polar cod (Benjamin J Laurel et al., 2019). Eye defects are also observed in bluefin tuna, yellowfin tuna, amberjack, Atlantic haddock, and more (Incardona et al., 2014, 2004; Meador and Nahrgang, 2019; Sørhus et al., 2016). Threespine stickleback size was not able to recover during a two-week depuration period, suggesting if stickleback can recover, like salmon, it would take longer than two weeks. As mentioned in 4.1, the reduction of snout-vent length and eye diameter could impair visual acuity and swim performance, thus impacting prey availability and predation and ultimately survival (Benjamin J Laurel et al., 2019).

The swim bladder is required for fish to rise in the water column, which is needed to capture prey and react to changes in the water environment. Crude oil can prevent swim bladders from inflating in zebrafish (Incardona et al., 2004; Li et al., 2019) resulting in a decrease in swim bladder area. In this study, threespine stickleback swim bladders area and length were not affected by crude oil exposure. Stickleback and zebrafish do have quite different swim bladder physiologies which may be a potential reason why crude oil exposure causes non-inflation of the swim bladder in zebrafish and not stickleback. Zebrafish are physostomes (fish with a pneumatic duct to the swim bladder) and stickleback are physoclists (fish without a pneumatic duct) (Price and Mager, 2020; Thomas and Ollevier, 1992). However, it is not known if threespine stickleback are transient physostomes or complete physoclists. Transient phystosomes are physo-clists that during the larval stage have a pneumatic duct to the swim bladder that allows for initial inflation of the swim bladder, but then atrophies after inflation.

### 4.3 Immune Gene Expression

PAH exposure upregulates both IL-1ß and TNF-α transcription associated with inflammation in other fish species (Bo et al., 2012; Nakayama et al., 2008; Song et al., 2012a, 2011; Volz et al., 2005). However, treatment alone did not have a significant effect on IL-1ß or TNF-α expression. Instead, there was a significant interaction of dpf and treatment on the expression of both, in which transcript levels of the genes tended to be lower over time in oil-treated fish and increased in the control fish. This is opposite to adult sheepshead minnow – where immune genes were downregulated after seven days of exposure to crude oil, but upregulated after 14 days of exposure (Jones et al., 2017).

Pathogen susceptibility and pathogen-induced mortality is increased in oil exposed fish (Arkoosh et al., 2002; Bayha et al., 2017; Song et al., 2011); thus, the decreased level of transcripts of both inflammatory genes over time may make threespine stickleback more susceptible to pathogens, increasing their mortality in situ. IL-1ß encodes for a cytokine that induces inflammation, fever, cell proliferation, differentiation, and pyroptotic cell death (Fink and Cookson, 2005; Martinon et al., 2002). The activation of IL-1ß can occur from sterile urban particulate matter, not just pathogens (Hirota et al., 2012). TNF-α encodes for a cytokine that induces inflammation, cell proliferation, apoptosis, lipid metabolism, and coagulation (Cryns et al., 1996; Szlosarek et al., 2006). TNF-α expression is also known to increase when exposed to aryl hydrocarbons receptor ligands like PAHs (Korashy and El-Kadi, 2006). However, the expression of both genes was not changed by oil exposure alone.

These genes are not the only inflammation markers that are influenced by crude oil. PAH exposure increases macrophage activity in some species (like killfish) but decreased activity in other species (like carp) (Reynaud and Deschaux, 2006). Splenic melano-macrophage centers also increase in number and size in sea trout and Gulf killfish when exposed to crude oil (Ali et al., 2014). Thus, future experiments in stickleback should include other markers of inflammation to determine if other inflammatory pathways are impacted by crude oil alone. In vivo experiments of threespine stickleback with combination oil and pathogen challenges, should also be performed to determine the potential pathogenic susceptibility due to immunotoxic effects of crude oil.

Genes not associated with inflammation should also be examined to determine whether crude oil affects immunity in stickleback in other ways (Arkoosh et al., 2002; Arkoosh and Collier, 2002; Bayha et al., 2017; Heintz et al., 2000). Primer sets for innate immune genes not associated with inflammation, such as the natural killer cell enhancing factor (NKEF-ß), are limited for threespine stickleback. NKEF-ß enhances the cytotoxicity of natural killer cells and protects cells against oxidative damage (Stutz et al., 2015). Nonspecific cytotoxic cells, which are similar to natural killer cells, are reduced in carp, oyster toadfish, and mummichog by PAH exposure (Reynaud and Deschaux, 2006). Thus, we hypothe-size that NKEF-ß could also be reduced by PAH exposure.

### 4.4 Potential Limitations

Fish oil toxicity studies typically expose subjects to high-energy water-accommodated fractions (HEWAFs) of crude oil (Incardona et al., 2015, 2014; Sørhus et al., 2016). HEWAFs are prepared by manual shaking of crude oil and water in a separatory funnel or by using a blender and blending the water and crude oil. The oil in this experiment was added to water and shaken on a benchtop orbital shaker before addition to the flasks. There was visible oil slicking and attachment to the sides of the tubes. The oil could potentially not have dispersed into the water as effectively as studies using HEWAFs. We were not able to perform analytical chemistry to confirm the concentration of oil as the volume of the water in the flasks was under the standard volume needed for gas chromatography mass spectrometry (1L). In the future, gas chromatography mass spectrometry could be used to determine ∑PAH.

## 5. Conclusions

Oil exposure in developing threespine stickleback has variable effects. Exposure does not impact the survival of developing stickleback, however exposure did significantly reduce eye diameter, which may impact visual acuity and swim performance and thus survival in the wild. Snout-vent length was also smaller in crude oil exposed stickleback, similar to salmon, which have reduced total length when exposed to oil (Heintz et al., 2000). The inflation of the stickleback swim bladder is not affected by oil exposure, unlike zebrafish (Incardona et al., 2004; Li et al., 2019). Oil exposure alone also does not affect the expression of innate immune genes, IL-1ß, or TNF-α. However, fish exposed to oil have a downregulation in the expression of both genes from 14 to 28 dpf, whereas fish not exposed to oil have an upregulation between time points. These results are contrary to other studies in which crude oil and PAHs upregulate genes associated with inflammation (Bo et al., 2012; Nakayama et al., 2008; Szlosarek et al., 2006; Volz et al., 2005). Thus, developing threespine stickleback respond to crude oil exposure in some similar manners to other fish (reduced growth and no changes in survival), but differently in other ways (swim bladder inflation and no upregulation of genes associated with inflammation). Of the traits impacted by crude oil exposure alone, recovery was not achieved in the two-week depuration period. The trajectory of innate immune gene expression was also altered in oil exposed fish as expression increased over time in control fish but decreased over time in oil exposed fish. Taken together these results suggest that threespine stickleback need more than two-weeks to recover from the toxic effects of crude oil.

## Acknowledgements

The authors thank Rachael Kramp, Levi Wegner, Sabrina Hock, Ryan Lucas, Chris Barahona, Copper Danner, Nadia Sherman, Catherine D’Amelio, and Anastastia Hanson for providing help with experimental set up, dissections, and fish husbandry, Patrick Tomco for providing the Alaska North Slope crude oil used for this experiment and chemistry advice, Jesse Weber for help with statistics, and Douglas Causey for further feedback on the work.

## Data Availability

All data and the R markdown file used for statistical analysis for this paper can be found at https://github.com/kellysue94/effects_of_crude_oil_on_juvenile_threespine_stickleback_020719.

## References

Ali, A.O., Hohn, C., Allen, P.J., Ford, L., Dail, M.B., Pruett, S., Petrie-Hanson, L., 2014. The effects of oil exposure on peripheral blood leukocytes and splenic melano-macrophage centers of Gulf of Mexico fishes. Mar. Pollut. Bull. 79, 87–93. https://doi.org/10.1016/j.marpolbul.2013.12.036

Arkoosh, M., Casillas, E., Clemons, E., Huffman, P., Kagley, A., Collier, T., Stein, J., 2002. Increased susceptibility of juvenile chinook salmon to infectious disease after exposure to chlorinated and aromatic compounds found in contaminated urban estuaries. Mar. Environ. Res. 50, 470–471. https://doi.org/10.1016/s0141-1136(00)00225-7

Arkoosh, M.R., Collier, T.K., 2002. Ecological risk assessment paradigm for salmon: Analyzing immune function to evaluate risk. Hum. Ecol. Risk Assess. 8, 265–276. https://doi.org/10.1080/20028091056908

Bayha, K.M., Ortell, N., Ryan, C.N., Griffitt, K.J., Krasnec, M., Sena, J., Ramaraj, T., Takeshita, R., Mayer, G.D., Schilkey, F., Griffitt, R.J., 2017. Crude oil impairs immune function and increases susceptibility to pathogenic bacteria in southern flounder. PLoS One 12, 1–22. https://doi.org/10.1371/journal.pone.0176559

Bio-Rad, 2013. CFX96 TouchTM, CFX96 Touch Deep WellTM, CFX ConnectTM, and CFX384 TouchTM Real-Time PCR Detection Systems - Instruction Manual. Bio-Rad Lab. Inc. 1–178.

Bo, J., Gopalakrishnan, S., Fan, D.-Q., Thilagam, H., Qu, H.-D., Zhang, N., Chen, F.-Y., Wang, K.-J., 2012. Benzo[a]pyrene Modulation of Acute Immunologic Responses in Red Sea Bream Pretreated with Lipopolysaccharide. Environ. Toxicol. 29, 517–525. https://doi.org/10.1002/tox

Brown-Peterson, N.J., Krasnec, M.O., Lay, C.R., Morris, J.M., Griffitt, R.J., 2017. Responsesofjuvenilesouthern flounder exposed to Deepwater Horizon oil-contaminated sediments. Environ. Toxicol. Chem. 36, 1067–1076. https://doi.org/10.1002/etc.3629

Brown-Peterson, N.J., Krasnec, M.O., Takeshita, R., Ryan, C.N., Griffitt, K.J., Lay, C.R., Mayer, G.D., Bayha, K.M., Hawkins, W.E., Lipton, I., Morris, J., Griffitt, R.J., 2015. A multiple endpoint analysis of the effects of chronic exposure to sediment contaminated with Deepwater Horizon oil on juvenile Southern flounder and their associated microbiomes. Aquat. Toxicol. 165, 197–209. https://doi.org/10.1016/j.aquatox.2015.06.001

Carls, M.G., Hose, J.E., Thomas, R.E., Rice, S.D., 2000. Exposure of pacific herring to weathered crude oil: Assessing effects on ova. Environ. Toxicol. Chem. 19, 1649–1659. https://doi.org/10.1002/etc.5620190624

Cryns, V.L., Bergeron, L., Zhu, H., Li, H., Yuan, J., 1996. Specific Cleavage of α-Fodrin during Fas-and Tumor Necrosis Factor-induced Apoptosis Is Mediated by an Interleukin-1ß-con-verting Enzyme / Ced-3 Protease Distinct from the Poly (ADP-ribose) Polymerase Protease *. J. Biol. Chem. 271, 31277–31282.

Delunardo, F.A.C., Silva, B.F. da, Paulino, M.G., Fernandes, M.N., Chippari-Gomes, A.R., 2013. Genotoxic and morphological damage in Hippocampus reidi exposed to crude oil. Ecotoxicol. Environ. Saf. 87, 1–9. https://doi.org/10.1016/j.ecoenv.2012.09.029

Edmunds, R.C., Gill, J.A., Baldwin, D.H., Linbo, T.L., French, B.L., Brown, T.L., Esbaugh, A.J., Mager, E.M., Stieglitz, J., Hoenig, R., Benetti, D., Grosell, M., Scholz, N.L., Incardona, J.P., 2015. Corresponding morphological and molecular indicators of crude oil toxicity to the developing hearts of mahi mahi. Sci. Rep. 5, 1–18. https://doi.org/10.1038/srep17326

Fink, S.L., Cookson, B.T., 2005. Apoptosis, pyroptosis, andnecrosis:Mechanisticdescriptionofdeadanddying eukaryotic cells. Infect. Immun. 73, 1907–1916. https://doi.org/10.1128/IAI.73.4.1907-1916.2005

Gard, R., Bottorff, R.L., 2014. Stickleback — Juvenile Sockeye Salmon Interactions, in: A History of Sockeye Salmon Research, Karluk River System, Alaska, 1880-2010. pp. 277–290. https://doi.org/10.7755/TMSPO.125

Glippa, O., Brutemark, A., Johnson, J., Spilling, K., 2017. Early Development of the Threespine Stickleback in Relation to Water pH. Front. Mar. Sci. 4, 1–8. https://doi.org/10.3389/fmars.2017.00427

Heintz, R. a., Rice, S.D., Wertheimer, A.C., Bradshaw, R.F., Thrower, F.P., Joyce, J.E., Short, J.W., 2000. Delayed effects on growth and marine survival of pink salmon Oncorhynchus gorbuscha after exposure to crude oil during embryonic development. Mar. Ecol. Prog. Ser. 208, 205–216. https://doi.org/10.3354/meps208205

Hibbeler, S., Scharsack, J.P., Becker, S., 2008. Housekeeping genes for quantitative expression studies in the three-spined stickleback Gasterosteus aculeatus. BMC Mol. Biol. 9, 1–10. https://doi.org/10.1186/1471-2199-9-18

Hirota, J.A., Hirota, S.A., Warner, S.M., Stefanowicz, D., Shaheen, F., Beck, P.L., MacDonald, J.A., Hackett, T.L., Sin, D.D., Van Eeden, S., Knight, D.A., 2012. The airway epithelium nucleotide-binding domain and leucine-rich repeat protein 3 inflammasome is activated by urban particulate matter. J. Allergy Clin. Immunol. 129, 1116–1125. e6. https://doi.org/10.1016/j.jaci.2011.11.033

Hook, S.E., Lampi, M.A., Febbo, E.J., Ward, J.A., Parkerton, T.F., 2010. Temporal patterns in the transcriptomic response of rainbow trout, Oncorhynchus mykiss, to crude oil. Aquat. Toxicol. 99, 320–329. https://doi.org/10.1016/j.aquatox.2010.05.011

Incardona, J.P., Carls, M.G., Holland, L., Linbo, T.L., Baldwin, D.H., Myers, M.S., Peck, K.A., Tagal, M., Rice, S.D., Scholz, N.L., 2015. Very low embryonic crude oil exposures cause lasting cardiac defects in salmon and herring. Sci. Rep. 5, 1–13. https://doi.org/10.1038/srep13499

Incardona, J.P., Collier, T.K., Scholz, N.L., 2004. Defects in cardiac function precede morphological abnormalities in fish embryos exposed to polycyclic aromatic hydrocarbons. Toxicol. Appl. Pharmacol. 196, 191–205. https://doi.org/10.1016/j.taap.2003.11.026

Incardona, J.P., Gardner, L.D., Linbo, T.L., Brown, T.L., Esbaugh, A.J., Mager, E.M., Stieglitz, J.D., French, B.L., Labenia, J.S., Laetz, C.A., Tagal, M., Sloan, C.A., Elizur, A., Benetti, D.D., Grosell, M., Block, B.A., Scholz, N.L., 2014. Deepwater Horizon crude oil impacts the developing hearts of large predatory pelagic fish. PNAS. https://doi.org/10.1073/pnas.1320950111

Jones, E.R., Martyniuk, C.J., Morris, J.M., Krasnec, M.O., Griffitt, R.J., 2017. Exposure to Deepwater Horizon oil and Corexit 9500 at low concentrations induces transcriptional changes and alters immune transcriptional pathways in sheepshead minnows. Comp. Biochem. Physiol. - Part D 23, 8–16. https://doi.org/10.1016/j.cbd.2017.05.001

Korashy, H., El-Kadi, A.O.S., 2006. The role of aryl hydrocarbon receptor in the pathogenesis of cardiovascular diseases, Drug Metabolism Reviews. Informa Healthcare. https://doi.org/10.1080/03602530600632063

Laurel, Benjamin J., Copeman, L.A., Iseri, P., Spencer, M.L., Hutchinson, G., Nordtug, T., Donald, C.E., Meier, S., Allan, S.E., Boyd, D.T., Ylitalo, G.M., Cameron, J.R., French, B.L., Linbo, T.L., Scholz, N.L., Incardona, J.P., 2019. Embryonic Crude Oil Exposure Impairs Growth and Lipid Allocation in a Keystone Arctic Forage Fish - Supplimental Information. iScience 19, 1101–1113. https://doi.org/10.1016/j.isci.2019.08.051

Laurel, Benjamin J, Louise, A., Tiffany, L., Nathaniel, L., John, P., Laurel, B.J., Copeman, L.A., Iseri, P., Spencer, M.L., Hutchinson, G., Nordtug, T., Donald, C.E., Meier, S., Allan, S.E., Boyd, D.T., Ylitalo, G.M., Cameron, J.R., French, B.L., Linbo, T.L., Scholz, N.L., Incardona, J.P., 2019. Embryonic Crude Oil Exposure Impairs Growth and Lipid Allocation in a Keystone Arctic Forage Fish Embryonic Crude Oil Exposure Impairs Growth and Lipid Allocation in a Keystone Arctic Forage Fish. iScience 19, 1101–1113. https://doi.org/10.1016/j.isci.2019.08.051

Leung, Y.F., Dowling, J.E., 2005. Gene expression profiling of zebrafish embryonic retina. Zebrafish 2, 269–283. https://doi.org/10.1089/zeb.2005.2.269

Li, X., Xiong, D., Ding, G., Fan, Y., Ma, X., Wang, C., Xiong, Y., Jiang, X., 2019. Exposure to water-accommodated fractions of two different crude oils alters morphology, cardiac function and swim bladder development in early-life stages of zebrafish. Chemosphere 235, 423–433. https://doi.org/10.1016/j.chemosphere.2019.06.199

Martinon, F., Burns, K., Tschopp, J., 2002. The Inflammasome: A molecular platform triggering activation of inflammatory caspases and processing of proIL-β. Mol. Cell 10, 417–426. https://doi.org/10.1016/S1097-2765(02)00599-3

Meador, J.P., Nahrgang, J., 2019. Characterizing Crude Oil Toxicity to Early-Life Stage Fish Based On a Complex Mixture : Are We Making Unsupported Assumptions ? Environ. Sci. Technol. A--M. https://doi.org/10.1021/acs.est.9b02889

Milinkovitch, T., Kanan, R., Thomas-Guyon, H., Floch Le, S., 2011. Effects of dispersed oil exposure on the bioaccumulation of polycyclic aromatic hydrocarbons and the mortality of juvenile Liza ramada. Sci. Total Environ. 409, 1643–1650. https://doi.org/10.1016/j.scitotenv.2011.01.009

Milligan-Myhre, K., Small, C.M., Mittge, E.K., Agarwal, M., Currey, M., Cresko, W.A., Guillemin, K., 2016. Innate immune responses to gut microbiota differ between oceanic and freshwater threespine stickleback populations. DMM Dis. Model. Mech. 9, 187–198. https://doi.org/10.1242/dmm.021881

Moles, A., Rice, S.D., Korn, S.I.D., 1979. Sensitivity of Alaskan Freshwater and Anadromous Fishes to Prudhoe Bay Crude Oil and Benzene, in: TransactionsoftheAmericanFisheriesSociety.pp.408–414.

Nakayama, K., Kitamura, S.I., Murakami, Y., Song, J.Y., Jung, S.J., Oh, M.J., Iwata, H., Tanabe, S., 2008. Toxicogenomic analysis of immune system-related genes in Japanese flounder (Paralichthys olivaceus) exposed to heavy oil. Mar. Pollut. Bull. 57, 445–452. https://doi.org/10.1016/j.marpolbul.2008.02.021

Palm, R.C., Powell, D.B., Skillman, A., Godtfredsen, K., 2003. Immunocompetence of juvenile chinook salmon against Listonella anguillarum following dietary exposure to polycyclic aromatic hydrocarbons. Environ. Toxicol. Chem. 22, 2986–2994. https://doi.org/10.1897/02-561

Pollino, C.A., Holdway, D.A., 2003. Hydrocarbon-induced changes to metabolic and detoxification enzymes of the Australian crimson-spotted rainbowfish (Melanotaenia fluviatilis). Environ. Toxicol. 18, 21–28. https://doi.org/10.1002/tox.10098

Price, E.R., Mager, E.M., 2020. The effects of exposure to crude oil or PAHs on fish swim bladder development and function. Comp. Biochem. Physiol. Part - C Toxicol. Pharmacol. 238. https://doi.org/10.1016/j.cbpc.2020.108853

Raimondo, S., Hemmer, B.L., Lilavois, C.R., Krzykwa, J., Almario, A., Awkerman, J.A., Barron, M.G., 2016. Effects of Louisiana Crude Oil on the Sheepshead Minnow (Cyprinodon Variegatus) During a Life-Cycle Exposure to Laboratory Oiled Sediment. Environ. Toxicol. 1627–1639. https://doi.org/10.1002/tox.22167

Reynaud, S., Deschaux, P., 2006. The effects of polycyclic aromatic hydrocarbons on the immune system of fish: A review. Aquat. Toxicol. 77, 229–238. https://doi.org/10.1016/j.aquatox.2005.10.018

Robertson, S., Bradley, J.E., Maccoll, A.D.C., 2016. Measuring the immune system of the three-spined stickleback - investigating natural variation by quantifying immune expression in the laboratory and the wild. Mol. Ecol. Resour. 16, 701–713. https://doi.org/10.1111/1755-0998.12497

Song, J.Y., Nakayama, K., Kokushi, E., Ito, K., Uno, S., Koyama, J., Rahman, M.H., Murakami, Y., Kitamura, S.I., 2012a. Effect of heavy oil exposure on antibacterial activity and expression of immune-related genes in Japanese flounder Paralichthys olivaceus. Environ. Toxicol. Chem. 31, 828–835. https://doi.org/10.1002/etc.1743

Song, J.Y., Nakayama, K., Murakami, Y., Kitamura, S.I., 2011. Heavy oil exposure induces high moralities in virus carrier Japanese flounder Paralichthys olivaceus. Mar. Pollut. Bull. 63, 362–365. https://doi.org/10.1016/j.marpolbul.2011.01.020

Song, J.Y., Ohta, S., Nakayama, K., Murakami, Y., Kitamura, S.I., 2012b. A time-course study of immune response in Japanese flounder Paralichthys olivaceus exposed to heavy oil. Environ. Sci. Pollut. Res. 19, 2300–2304. https://doi.org/10.1007/s11356-012-0737-z

Sørhus, E., Incardona, J.P., Karlsen, Ø., Linbo, T., Sørensen, L., Nordtug, T., Van Der Meeren, T., Thorsen, A., Thorbjørnsen, M., Jentoft, S., Edvardsen, R.B., Meier, S., 2016. Crude oil exposures reveal roles for intracellular calcium cycling in haddock craniofacial and cardiac development. Sci. Rep. 6. https://doi.org/10.1038/srep31058

Stutz, W.E., Schmerer, M., Coates, J.L., Bolnick, D.I., 2015. Among-lake reciprocal transplants induce convergent expression of immune genes in threespine stickleback. Mol. Ecol. 24, 4629–4646. https://doi.org/10.1111/mec.13295

Szlosarek, P., Charles, K.A., Balkwill, F.R., 2006. Tumour necrosis factor-α as a tumour promoter. Eur. J. Cancer 42, 745–750. https://doi.org/10.1016/j.ejca.2006.01.012

Thomas, K., Ollevier, F., 1992. Paratenic hosts of the swimbladder nematode Anguillicola crassus. Dis. Aquat. Organ. 13, 165–174. https://doi.org/10.3354/dao013165

Thorn, M.W., Morbey, Y.E., 2018. Egg size and the adaptive capacity of early life history traits in Chinook salmon (Oncorhynchus tshawytscha). Evol. Appl. 11, 205–219. https://doi.org/10.1111/eva.12531

Volz, D.C., Bencic, D.C., Hinton, D.E., Law, M.J., Kullman, S.W., 2005. 2,3,7,8-Tetrachlorodibenzo-p-dioxin (TCDD) induces organ-specific differential gene expression in male Japanese medaka (Oryzias latipes). Toxicol. Sci. 85, 572–584. https://doi.org/10.1093/toxsci/kfi109

